# Muscle Fibroblasts and Stem Cells Stimulate Motor Neurons in An Age and Exercise-Dependent Manner

**DOI:** 10.1101/2024.08.19.608387

**Authors:** Casper Soendenbroe, Peter Schjerling, Cecilie J.L. Bechshøft, Rene B. Svensson, Laurent Schaeffer, Michael Kjaer, Bénédicte Chazaud, Arnaud Jacquier, Abigail L. Mackey

**Author notes:** Co-correspondence.

## Abstract

Exercise preserves neuromuscular function in ageing through unknown mechanisms. Skeletal muscle fibroblasts (FIB) and stem cells (MuSC) are abundant in skeletal muscle and reside close to neuromuscular junctions, but their relative roles in motor neuron maintenance remain undescribed. Using direct co-cultures of embryonic rat motor neurons with either human MuSC or FIB, RNA sequencing revealed profound differential regulation of the motor neuron transcriptome, with FIB generally favoring neuron growth and cell migration and MuSC favoring production of ribosomes and translational machinery. Conditioned medium from FIB was superior to MuSC in preserving motor neurons and increasing their maturity. Lastly, we established the importance of donor age and exercise status and found an age-related distortion of motor neuron and muscle cell interaction that was fully mitigated by lifelong physical activity. In conclusion, we show that human muscle FIB and MuSC synergistically stimulate the growth and viability of motor neurons, which is further amplified by regular exercise.

## Introduction

The number of alpha motor neurons in the spinal cord declines with age (*1*, *2*), leading to muscle fiber denervation (*3*), that if not resolved, causes irreversible loss of muscle fibers (*4*, *5*). Lost muscle fibers are replaced by fibrotic tissue (*6*), driven by activation of the mesenchymal fibroblast cells (*7*, *8*). Muscle fibroblasts (FIB) reside between muscle fibers, constitute ≈10-15% of all nuclei in muscle (*9*), and are necessary for efficient recovery following muscle injury (*10*, *11*). Upon denervation, fibroblasts are stimulated to proliferate, especially in junctional regions (*12*), and it has long been speculated that they might facilitate nerve outgrowth by producing adhesive molecules, such as Neural-Cell-Adhesion-Molecule and Fibronectin (*8*), remodel the extracellular matrix (*13*), or secrete neurotrophic factors (*14*). Moreover, depletion of PDGFRα+ positive cells, a marker of the mesenchymal fibro-adipogenic progenitor cell, that gives rise to FIB, leads to more partial and fully denervated neuromuscular junctions (NMJ) (*15*), hinting at a role for FIB in securing and maintaining muscle innervation. Muscle stem cells (MuSC) play a pivotal role in regeneration following muscle injury (*16*), and recent studies have highlighted substantial cell heterogeneity (*17*) and involvement in a multitude of cellular processes (*18*), including maintaining muscle innervation (*19*). *In vitro*, MuSC communicate with motor neurons directly through retrograde signaling (*20*) and indirectly through secretion of circulating factors (*21*). Yet, the relative importance of FIB and MuSC interaction with motor neurons, through direct and indirect means, has not been studied. Although still debated (*22*, *23*), the decay of motor neurons is believed to begin during middle age leading to an often temporary denervation of muscle fibers (*1*), as they are rescued by collateral reinnervation by closely located axon terminals (*24*). By electromyography, this presents as larger and more complex motor units (*25*). Later in life, the decay of motor neurons accelerates and/or the efficiency of reinnervation declines, leading to permanent denervation of muscle fibers (*26*). Both animal and human studies have indicated that exercise mitigates some of the detrimental age-effects on the neuromuscular system (*27*). In addition to superior physical function, master athletes show attenuated signs of denervation and larger motor units (*28*, *29*), while heavy resistance exercise reduces muscle markers of denervation in older individuals (*30*), suggesting retained plasticity in the neuromuscular system of aged individuals. Importantly, the protective effect of exercise is partly driven by improving the ability to reinnervate denervated muscle fibers, whereas the effect of exercise on the preservation of motor neurons remains unexplored.

Here, we tested the hypothesis that FIB, in addition to MuSC, interact with motor neurons through both direct and indirect means. We found that the motor neuron transcriptome was profoundly altered when directly exposed to FIB compared to MuSC. FIB upregulated pathways related to neuron growth and cell migration, highlighting an important role in stimulating motor neurons. Motor neurons cultured in conditioned medium from FIB showed overall higher survival, and higher expression of the mature motor neuron marker, Choline Acetyltransferase (ChAT), than when cultured in medium from MuSC. Finally, we investigated the effect of donor age and exercise status upon the motor neuron transcriptome. Lifelong exercisers were phenotypically different from sedentary individuals, with higher muscle function and lower circulating levels of the neuromuscular disturbance biomarker, C-terminal Agrin Fragment. *In vitro*, this effect translated into higher survival of motor neurons when exposed to FIB and MuSC from exercisers compared to sedentary individuals. Together, our findings highlight that motor neurons are positively influenced by cells from lifelong exercisers, and that FIB in particular may be a key player in maintaining neuromuscular function in aging.

## Results

### Differential regulation of motor neuron gene expression by muscle stem cells and fibroblasts

Fibroblasts reside in the interstitial space in muscle and are often observed close to NMJs (*15*). To investigate if and how fibroblasts influence the motor neuron transcriptome, a direct cell culture model was established, using primary motor neurons isolated and purified from E14 rat embryos (*31*), and primary human MuSC and FIB isolated and purified from human muscle biopsies (*32*). The compatibility of human muscle cells with rat motor neurons was confirmed by generating highly differentiated myotubes that contracted only when cultured with spinal cord explants (Movie S1-S3). Species-specific RNA sequencing was analyzed for 14 male participants, containing highly pure fractions of both MuSC and FIB (Fig. 1.A). The participants represented were young (n=4, Young), old sedentary (n=4, SED) and old lifelong exercisers (n=6, LLEX). MuSC or FIB were cultured alone for 24 hours, after which motor neurons were added, and the cultures continued for another 24 hours (Fig. 1.B). FIB was clearly different from MuSC, with 6968 and 6699 human genes upregulated or downregulated in MuSC compared to FIB, respectively, corresponding to 27% of all detected genes (Fig.1.C). Both FIB and MuSC expressed several well-known canonical or cell-type specific markers such as *PDGFR*α (FIB) and *MYH7*, *MYH2*, *CASQ2*, *PAX7*, *MYL2* and *TNNC1* (MuSC). Additionally, FIB expressed markers that, using single-cell RNA sequencing, have been used to identify specific mesenchymal cell subpopulations (*33*, *34*), such as *DCN*, *CD34* and *COL6a3*.

**Fig. 1.**
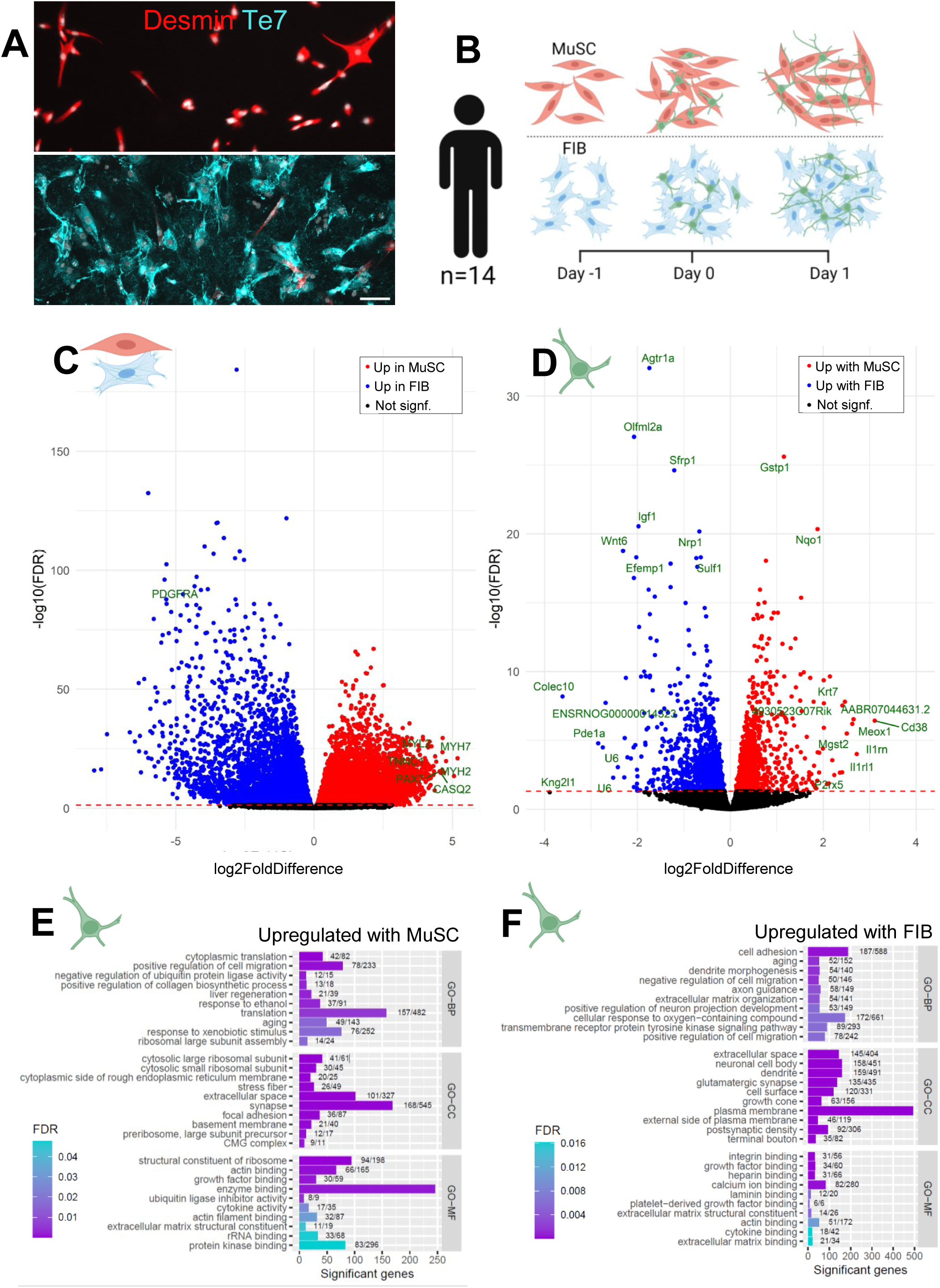
Differential regulation of motor neuron gene expression by muscle stem cells and fibroblasts. **A**) Representative image of MuSC (red=desmin) and FIB (blue=Te7) in vitro. Scale bar, 100 μm. **B**) Experimental setup of direct co-culture experiments: E14 rat motor neurons were plated on human primary MuSC or FIB for 1 day and analyzed by RNA sequencing. **C**) Volcano plot showing differentially regulated human MuSC vs FIB genes. Blue and red are upregulated in FIB and MuSC, respectively. Cell type specific markers are highlighted. **D**) Volcano plot showing differentially regulated rat genes upon exposure to human MuSC vs FIB. Blue and red are upregulated with FIB and MuSC, respectively. Top 10 lowest FDR, and Top 10 highest log2FoldDifference for each cell type is highlighted. **E**) Pathway analysis showing top-10 GO-BP, GO-CC and GO-MF terms upregulated in motor neurons exposed to MuSC vs FIB. **F**) Pathway analysis showing top-10 GO-BP, GO-CC and GO-MF terms upregulated in motor neurons exposed to FIB vs MuSC. Level of significance is indicated by color. Number of differentially regulated genes and total number of genes is indicated for each term. **C-F**) Human n: Paired MuSC and FIB from 14 individuals. Rat n: Embryos from 7 dams. Statistics: Data were analyzed with DESeq2 and TopGO (elim), see bioinformatics section. Abbreviations: MuSC, muscle stem cell; FIB, muscle fibroblast.

For motor neurons (rat genome), it was observed that 1288 and 1285 genes were upregulated or downregulated when cultured with MuSC compared to FIB, respectively, corresponding to 11% of all detected genes (Fig.1.D). These included genes involved in many different processes, such as *Olfml2*α, involved in extracellular matrix binding activity, *Wnt6*, involved in secretion of signaling proteins and *Cd38*, involved in intracellular calcium metabolism.

To probe whether key neuronal processes were affected by cell type, pathway analyses were conducted using Gene Ontology (GO-terms) (Fig.1.E-F). In all cases, the top 10 pathways are shown. Within the GO Biological Processes (GO-BP) domain, MuSC mostly increased pathways associated with ribosomes and translation, whereas FIB increased several pathways associated with neurons and their projection of neurites. Similarly, within the GO Cellular Components (GO-CC) domain, MuSC increased a pathway associated with “synapse”, and other pathways, such as “focal adhesion”, “extracellular space” and “basement membrane”. In contrast, FIB increased several pathways associated with neurite growth, such as “terminal bouton”, “dendrite” and “neuronal cell body”. Motor neuron specific genes were identified from two single-cell RNA sequencing data sets of murine spinal cord cells and the regulation of these genes in the present data set was investigated (Fig.2.A-B). In a study by Delile et al., analyzing spinal cord cells during different stages of development (*35*), 38 genes specifically expressed in motor neurons were identified (Fig.2.A). A second study, by Blum et al., identified 33 genes specific to motor neurons in adult mice (*36*) (Fig.2.B). The 71 extracted genes included established motor neuron markers, such as *Ret* (embryonic) and *Chat* (mature), but also genes not previously associated with motor neurons. Interestingly, in our sequencing data, 23 out of 25 differentially expressed genes (DEGs) showed higher expression in the rat cells exposed to FIB compared to MuSC. These genes include Ret and Chat, but also less established marker genes such as Slc5a7, Pcdh7 and Megf11. The genes favored by MuSC compared to FIB were *Nr2f2* and *Ebf2*, two transcription factors involved in several developmental processes. Of special note, two genes involved in chemical synapse transmission, *Sv2c* and *Slc5a7*, and one gene in neurogenesis, *Tubb3*, were all upregulated in FIB compared to MuSC.

**Fig. 2.**
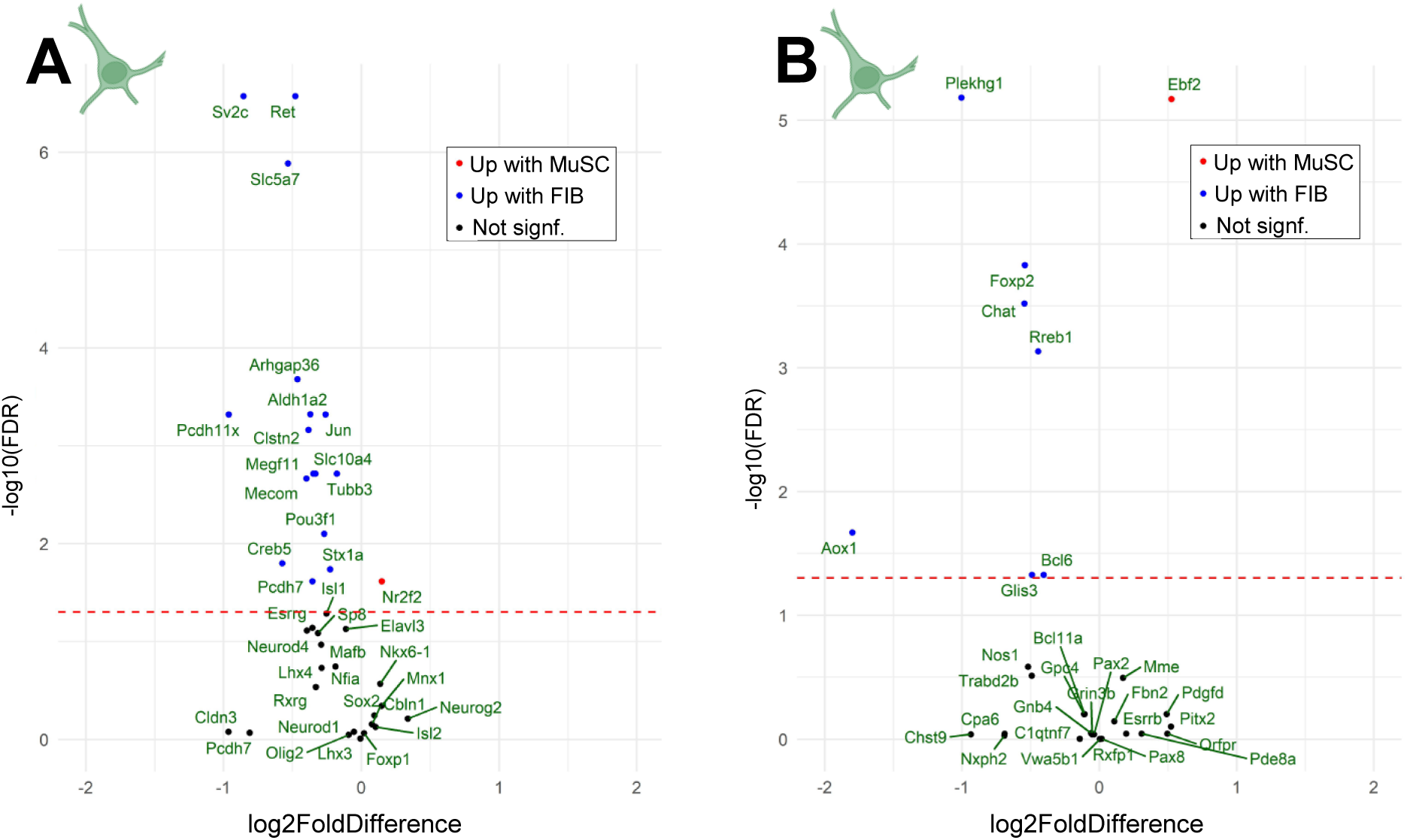
Motor neuron specific genes are upregulated by fibroblasts. Motor neuron specific genes identified from two single-cell RNA sequencing data sets of murine spinal cord cells. **A**) Volcano plot showing differentially regulated rat motor neuron specific genes from reference 34 upon exposure to human MuSC vs FIB. Blue and red are upregulated with FIB and MuSC, respectively. **B**) Volcano plot showing differentially regulated rat motor neuron specific genes from reference 35 upon exposure to human MuSC vs FIB. Blue and red are upregulated with FIB and MuSC, respectively. Human n: Paired MuSC and FIB from 14 individuals. Rat n: Embryos from 7 dams. Statistics: Data were analyzed with DESeq2, see bioinformatics section. Abbreviations: MuSC, muscle stem cell; FIB, muscle fibroblast.

### Fibroblast conditioned medium preserves motor neurons and increases their maturity

Next, we wanted to explore whether conditioned medium from FIB or MuSC, containing all secreted factors, evoked changes in motor neuron growth and viability (Fig.3.A). To do that, primary muscle cells were isolated from muscle biopsies collected from 10 younger and 10 older female participants. The cells were expanded in culture, sorted into highly pure fractions of FIB or MuSC and allowed to differentiate for 7 days, during which conditioned medium was collected. The younger and older participants were phenotypically different, as evidenced by differences in muscle strength and muscle fiber size (Fig.3.B). Furthermore, as previously reported, the MuSC of older participants demonstrated impaired fusion capacity compared to younger participants (*37*). Conditioned medium from FIB and MuSC was added to motor neurons cultures for 1 and 2 days, and the cells were analyzed by immunofluorescence and RNA sequencing.

**Fig. 3.**
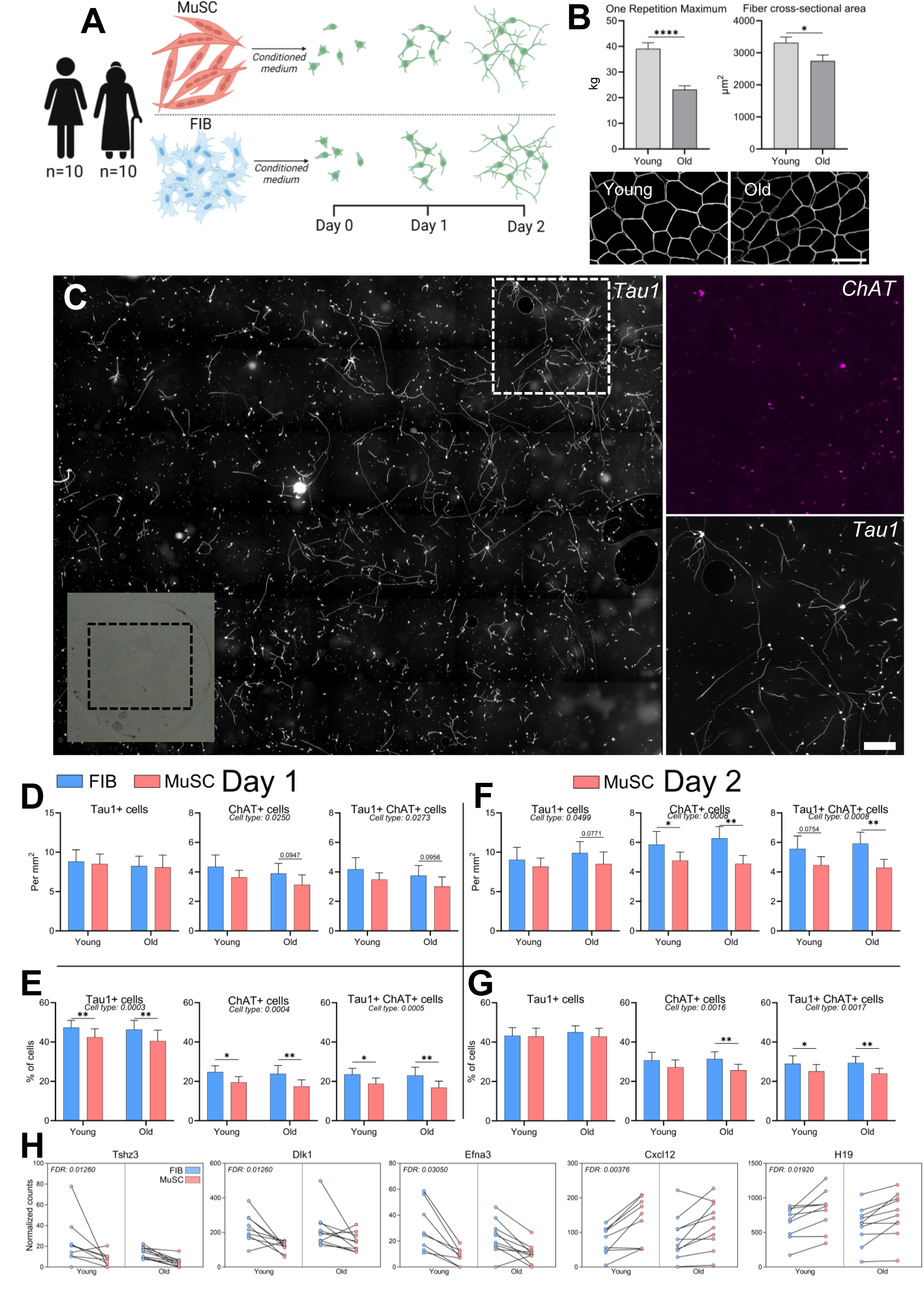
Fibroblast conditioned medium preserves motor neurons and increases their maturity. **A)** Experimental setup of conditioned medium experiments: E14 rat motor neurons were plated in Poly-L-Ornithine and laminin coated wells, in 40% conditioned medium from MuSC or FIB, and analysed after 1 and 2 days by immunocytochemistry and RNA sequencing. **B**) One-repetition maximum and muscle fiber cross-sectional area. Data are means ± SEM. Statistics: Data were analyzed by unpaired two-tailed t-tests.* indicates p<0.05, **** p<0.0001. Representative image muscle cross-section immunofluorescently stained with dystrophin. Scale bar, 100 μm. **C**) Example of entire coverslip used for analysis. Insert shows zoomed-in area. Scale bar, 250 μm. Magenta=ChAT, grey=Tau1. **D-G**) Immunocytochemical analyses of Tau1+ and ChAT+ cells, either per area (D,F), or as a percentage of all cells (E,G), after 1 (D,E) and 2 (F,G) days. Data are means ± SEM. Human n: Old: 10, Young: 10. Rat n: 7. Statistics: Data were analyzed by two-way repeated measures ANOVA (age group x cell type), with Fisher’s LSD posthoc test. Main effects are written and post hoc tests indicated by *p<0.05, **p<0.01. **H**) Differentially regulated rat genes upon exposure to human MuSC vs FIB after 1 day, displayed as paired values. Blue and red is FIB and MuSC, respectively. N: Old: 10, Young: 9. Rat n: 7. Data were analyzed with DESeq2, see bioinformatics section. Abbreviations: MuSC, muscle stem cell; FIB, muscle fibroblast.

While there was no effect of age, exposing motor neurons to conditioned medium from FIB for 1 day increased the number of ChAT^+^ and Tau1^+^ChAT^+^ motor neurons per mm^2^, while also increasing the percentages of Tau1^+^ neurons, and ChAT^+^ and Tau1^+^ChAT^+^ motor neurons compared to MuSC conditioned medium (Fig.3.D-E). The latter cell type differences were observed in both age groups. Similarly, after 2 days of culture, where neurons continued to thrive, a similar pattern for superior support from FIB vs MuSC was observed and was even more substantial (Fig.3.F-G).

We then continued to explore differences in the rat cells at the gene expression level and found two upregulated (*Cxcl12* and *H19*) and three downregulated (*Tshz3*, *Dlk1* and *Efna3*) genes with conditioned medium from MuSC compared to FIB, respectively, corresponding to 0.02% of the total gene pool to be influenced (Fig.3.H). Of these five DEGs, *Efna3* and *Cxcl12* appear to have the most relevance to neuronal processes, as *Efna3* is involved in axon guidance (*38*) and *Cxcl12* is known to promote NMJ regeneration and axonal extension (*39*). We observed that *Efna3* was higher in FIB compared to MuSC, and *Cxcl12* was higher in MuSC compared to FIB, suggesting that FIB and MuSC have complementary effects on motor neurons. *Tshz3*, *Dlk1* and *H19* have also been linked to various neuronal processes (*40–42*). Of note, all five DEGs were also detected in the direct co-culture experiments, where two of them (*tshz3* and *Dlk1*) were similar between FIB and MuSC conditions, *Efna3* was higher in FIB compared to MuSC, and both *Cxcl12* and *H19* were higher in MuSC compared to FIB.

### Sedentary ageing is associated with reduced muscle performance and neuromuscular disturbance

Next, we wanted to explore whether lifelong recreational exercise confers protective effects on motor neurons. To that end, men who had been recreationally active for most of their life were compared with a sedentary control group of similar age, as well as with a young sedentary control group. These individuals were part of a larger study, in which we have previously shown that lifelong exercise is associated with preserved muscle content of MuSCs and better muscle innervation status (*43*). Despite leg lean mass being similar between groups, isometric maximum voluntary contraction (MVC) strength was 39 and 26% lower in the old SED and LLEX, respectively, compared to Young (Fig. 4.B-C). This in turn translated into a 33 and 21% difference between Young and SED and LLEX, respectively, in muscle quality, as assessed by the specific force index (Fig. 4.D). The 7 participants in LLEX had vastly different training backgrounds, not all performing activities requiring maximal force output. As such, we wanted to evaluate their capacity to repeatedly perform maximal force output under fatiguing conditions, as this more closely resembled their normal activities (resistance exercise, football, cycling, rowing, gymnastics). This was evaluated as a muscle performance index, reflecting an average of the% of MVC sustained during the first, fifth and tenth repetition performed during eight sets of concentric knee extensions. As expected, LLEX performed 14% better than SED, but surprisingly, LLEX were also 12% better than Young participants (Fig. 4.E), showing that LLEX were accustomed to exhausting exercise. The immunohistological assessment of the muscle biopsies revealed that muscle fiber size was 39% smaller in SED compared to Young, while no significant difference was observed between LLEX and Young (p=0.08) (Fig.4.G). Lastly, to probe whether these functional and structural differences could be sourced to the neuromuscular system, serum concentration of C-terminal Agrin Fragment (CAF), a biomarker indicative of neurological impairment was measured. As expected, CAF was elevated in SED compared to both Young and LLEX, while no difference was observed between Young and LLEX (Fig. 4.F). Together, these data demonstrate clear phenotypic differences of being sedentary or being modestly physically active, throughout life.

**Fig. 4.**
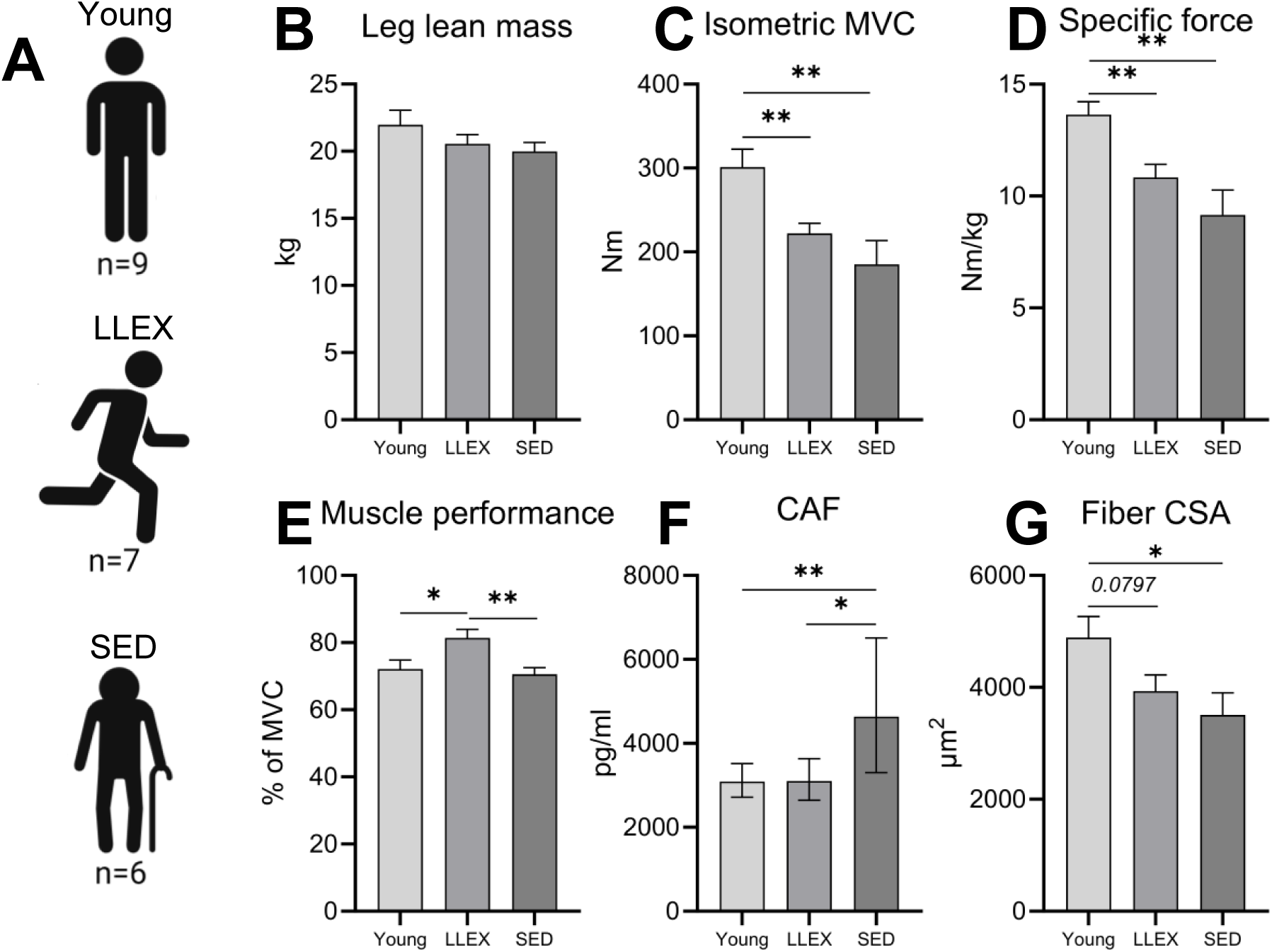
Sedentary ageing is associated with reduced muscle performance and neuromuscular disturbance *in vivo*. **A**) In vivo characterization of male participants. **B**) Leg lean mass, measured by DEXA. **C**) Isometric unilateral knee extension MVC, measured in a dynamometer. **D**) Specific force, calculated as MVC per leg lean mass. **E**) Muscle performance, measured as force exerted during repeated maximal knee extension concentric contractions, expressed relative to MVC. **F**) CAF, measured in plasma by ELISA. Statistical analysis was conducted on log transformed values. **G**) Muscle fiber cross sectional area measured using immunohistochemical analyses of muscle biopsy cross-sections. All data are means ± SEM except CAF which is shown as geometric means with 95% CI. N: LLEX: 7, SED: 6, Young: 9. Statistics: Data were analyzed by unpaired two-tailed t-tests, with significance indicated by *p<0.05, **p<0.01, ***p<0.001. Abbreviations: DEXA, dual-energy X-ray absorptiometry; MVC, maximal voluntary contraction; CAF, C-terminal Agrin Fragment

### Lifelong exercise facilitates motor neuron survival *in vitro*

Using the same experiment as in figure 1, but now focusing on group rather than cell type differences, we aimed to investigate the influence of donor age and exercise background, on motor neuron survival and neurite growth (Fig.5.A-B). Motor neurons were co-cultured with either FIB or MuSC for 1 and 2 days, and then processed for immunocytochemistry and RNA sequencing. The immunocytochemical analyses revealed that 53% more motor neurons survived when exposed to MuSC from LLEX compared to SED, with no significant difference between LLEX and Young (Fig.5.C). This finding was mirrored in FIB conditions (Fig.5.D). In both FIB and MuSC conditions, no differences between groups were observed for number or length of neurites.

**Fig. 5.**
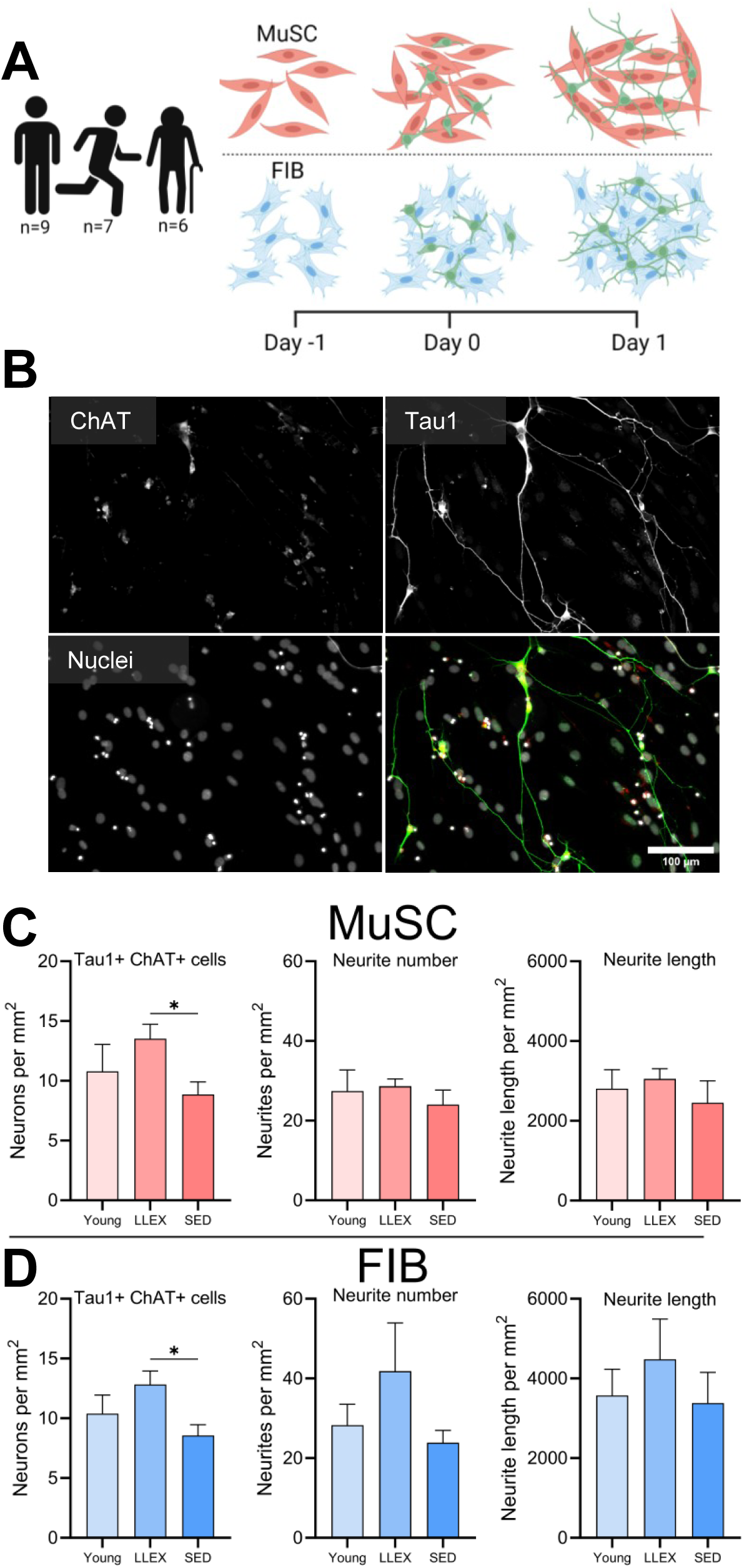
Lifelong exercise facilitates motor neuron survival *in vitro*. **A**) Experimental setup of direct co-culture experiments: Motor neurons were plated on MuSC and FIB isolated from muscles of Young, LLEX and SED, for 1 day and analyzed by immunocytochemistry and RNA sequencing. **B**) Representative image co-cultures stained with ChAT (red), Tau1 (green) and nuclei (grey). Scale bar, 100 μm. **C-D**) Number of Tau1+ and ChAT+ cells, number of neurites and length of neurites in MuSC (C) and FIB (D) conditions. Data are shown as means ± SEM. Human n (MuSC): LLEX: 7, SED: 6, Young: 9. Human n (FIB): LLEX: 5, SED: 4, Young: 6. Rat n: 7. Statistics: Data were analyzed by unpaired two-tailed t-tests, with significance indicated by *p<0.05. Abbreviations: MuSC, muscle stem cell; FIB, muscle fibroblast.

At the gene expression level, after FDR adjustment, there were 6 and 2 DEGs in MuSC and FIB, respectively, when comparing Young with Old (LLEX and SED combined) (Fig. S1). *PCDH10*, *CDH8*, *GABRB2* and *ENSG00000236536* were higher in MuSC from Old compared to Young, while *NEFH* and *TINAGL1* were higher in Young compared to Old. In FIB, *TRHDE-AS1* was higher in Old and *RANBP17* was higher in Young.

## Discussion

Fibroblasts (FIBs), and their precursors, the fibro-adipogenic progenitors, are abundant interstitial cells in skeletal muscle (*9*), that are required for maintaining tissue homeostasis and supporting regeneration following injury (*44*). Recent studies highlight a role for FIB in facilitating and maintaining muscle innervation, accomplished indirectly through interactions with Schwann cells (*15*), glial cells (*13*) and regulatory T cells (*45*). However, an incomplete understanding of how FIB interacts directly with motor neurons precludes targeted molecular interventions. We have now demonstrated that motor neuron gene expression is profoundly altered when exposed directly to FIBs compared to muscle stem cells (MuSC); established that conditioned medium of FIBs is superior to MuSC in promoting motor neuron maturation and neurite growth; and showed that survival of motor neurons is influenced by age and exercise status of muscle cell donors. These findings raise several points of interest.

The interaction of motor neurons with other cell types is predominantly focused on MuSC (*46*), likely owing to their importance in donating myonuclei to the muscle fiber itself (*18*). Yet, a recent focus on other, non-myogenic cells, such as FIB and fibro-adipogenic progenitors (*14*), has shown interactions between them and motor neurons, albeit mediated through tertiary cells (*13*, *15*, *45*). In the present study, motor neuron transcriptome was profoundly altered when exposed to FIB compared to MuSC, showing that FIB directly interacts with motor neurons through cell-to-cell contact. A high abundance of upregulated pathways associated with neurite growth further posits FIB as a key regular of neuritogenesis, which, in addition to their well-known remodeling of extracellular matrix (*14*), highlights a key role in establishing and maintaining muscle innervation. This extends the findings of Uezumi et al., who showed, in mice, that depletion of fibro-adipogenic progenitors increased the number of partially and fully denervated NMJs (*15*). Through subsequent *in vitro* experiments they observed a signaling cascade going from fibro-adipogenic progenitors through Schwann cells, which encapsulate NMJs and intramuscular nerves (*47*), to motor neurons. Our findings promote FIB to being a cell that, equivalent to MuSC (*20*), directly stimulates neuritogenesis, which again underscores that maintenance of muscle innervation is a synergistic endeavor involving multiple cell types.

Overall, the arrangement that FIB, in synergy with MuSCs, exert a direct, non-mediated effect on motor neurons is feasible. This is so, as there is a close anatomical proximity of these cells to both NMJs and intramuscular nerves (*15*), through which factors can be taken up and transported retrogradely to the motor neuron in the spinal cord (*48*). Further, it can be speculated that FIB, similar to MuSC, secrete molecules that are taken up in capillaries, enter general circulation, and exert endocrine functions in other tissues (*49*). Crucially, the exact origins of most secreted factors from skeletal muscle, of which there are hundreds or even thousands (*50*), remain unclear. It is however clear that muscle derived factors can be both neuro-hostile or neuro-supportive in nature (*51*), which is further underscored by studies with Amyotrophic Lateral Sclerosis muscle cells which secrete neuro-toxic vesicles (*52*). As such, the beneficial effect of FIB conditioned medium on motor neuron maturity and viability, in direct comparison with MuSC conditioned medium, observed in the present study, suggests that FIBs from healthy young and older adults, in general, is a source of neuro supportive factors.

MuSCs *in vitro*, retain key *in vivo* properties (*53*), such as improved lipid oxidation in trained compared to sedentary individuals (*54*) and growth capacity in older compared to younger individuals (*37*). Similarly, surgical denervation through nerve transection, which is an extreme model of disuse, increases the number, and alters the transcriptomic profile and *in vitro* properties of fibro-adipogenic progenitors (*6*, *13*). In our cohort of male participants, lifelong exercisers were phenotypically different from their sedentary peers in terms of muscle function and neuromuscular properties. These phenotypic hallmarks were imprinted on both FIB and MuSC *in vitro*, making them more neuro-supportive. This finding has important implications for exercise recommendations for the general population. While still debated (*55*), there is substantial evidence from both human cadaver studies (*1*, *56*, *57*) and electrophysiological investigations (*2*), that the number of motor neurons, or by proxy large-diameter myelinated axons, decline from around the age of 60 to 70 and is lower in older compared to younger individuals. This irreversible age-related loss of motor neurons causes muscle fiber denervation (*58*), and denervated muscle fibers atrophy and disappear, unless they are reinnervated by adjacent neurites (*3*). It is believed that for most of life, the capacity to reinnervate is well preserved and clearly outweighs the need. As such, older trained runners have larger motor units and preserved muscle mass compared to non-runners (*28*, *59*), yet there is clearly also an upper limit for how many muscle fibers each motor neuron can control (*60*). So while exercise can, even in advanced age, stimulate muscle fiber reinnervation (*30*), another, equally important parameter for healthy ageing is the preservation of motor neurons themselves. The finding herein that both FIB and MuSC from lifelong exercisers preserve motor neurons *in vitro*, suggests, that for optimal preservation of muscle function with ageing, exercise should be initiated prior to the beginning of motor neuron decay. This, in turn, also amplifies the capacity for reinnervation in later stages of ageing, as the pool of motor neurons is larger.

This study has limitations and there are outstanding questions for future research. Different types of exercise lead to specific physiological adaptations (*61*), and since the lifelong exercisers included in the present study performed many different types of exercise, it is not possible to say which type of exercise is superior for protecting motor neurons. Furthermore, while having access to adult human motor neurons would be beneficial, it is however not possible to isolate sufficient amounts of cells from live (or deceased) humans, necessitating the use of either inducible pluripotent or embryonic stem cells, which has other limitations (*62*).

In conclusion, FIB, MuSC and motor neurons represent a mutually supporting triad, where cells interact to maintain neuromuscular homeostasis through both direct and indirect means. This occurs in an age and exercise dependent manner, meaning, that while taking up exercise at any age will provide many health benefits, there may be a point of no return when it comes to motor neuron preservation. Optimal preservation of motor neurons may require maintaining an active lifestyle throughout life.

## Materials and Methods

### Experimental Design

The study was designed to compare whether and how muscle fibroblasts and muscle stem cells interact with motor neurons. Moreover, we aimed to investigate whether muscle cells, isolated from lifelong exercisers, exerted a protective and supportive effect on motor neurons. These objectives were investigated *in vitro* using primary muscle cells from human donors and primary motor neurons from rat embryos and were analyzed by immunocytochemistry and species-specific RNA sequencing. A short period (24-48 hours) of co-culture was chosen for the experiments, as differences at the gene expression level were expected to be greater initially. All analyses were conducted under blinded conditions (donor, cell type, group). All sample sizes, representing biological replicates, are reported in the material and methods section and in the appropriate figure legends.

### Human participants

Human procedures were approved by The Committees on Health Research Ethics for The Capital Region of Denmark (Ref: H-19000881, H-15017223) and were conducted according to the standards set by the Declaration of Helsinki. For this study, isolated muscle cells from 12 younger and 11 older females, as well as 9 younger and 13 older males, were obtained in relation to two previous studies (*37*, *43*). Of the 13 older males, 7 were life-long exercisers, having performed various types of physical activity (endurance, strength and mixed), non-competitively for most of their lives. Age, height, weight, and BMI were determined for all participants (Table S1). For the females, 1-repetition maximum strength was determined in unilateral leg extension exercise (details in reference 36). For the males, leg lean mass was determined by dual-energy X-ray absorptiometry (DEXA), maximal voluntary contraction (MVC) was measured in dynamometer, and a muscle performance index was determined by measuring force output during repeated maximal knee extension concentric contractions (details in reference 42). Venipuncture blood samples were collected from the males and vastus lateralis muscle biopsies were collected from all participants, using the Bergström biopsy needle technique (*63*). A part of the sample was embedded for histology and frozen in liquid nitrogen-cooled isopentane. The remaining tissue was used to isolate cells for *in vitro* experiments.

### Muscle cell isolation and sorting

The cell isolation has previously been described in detail (*37*, *43*). Briefly, muscle tissue was enzymatically and mechanically digested, and the released cells were then cultured in growth medium (C-23060; PromoCell) with 1% L-glutamine-penicillin-streptomycin solution (G6784; Sigma) and 15% fetal bovine serum (FBS; ALB-S1810, Biowest) for 6-9 days with medium change every 2-3 days. When confluent, cells underwent magnetic-activated cell sorting using a CD56 antibody (130-050-401; Miltenyi Biotec), producing a CD56+ fraction consisting of muscle stem cells (MuSC) and a CD56-fraction consisting of muscle fibroblasts (FIB) (*32*). Sorted cells were frozen until used for experiments.

### Animals

The use and care of rats in this study were carried out in accordance with the law on animal experiments in Denmark (Law on Animal Experiments in Denmark, LBK nr. 63, 19 January 2024) and Directive 2010/63/EU with the license number 2017-15-0201-01364) from the Animal Inspectorate, Ministry of Food, Agriculture and Fisheries, Denmark. No experiments were performed on live animals. A total of 14 timed-pregnant Sprague-Dawley rats, that were purchased from Taconic Biosciences (Ejby, Denmark), arrived at the research facility, one week prior to the experimental procedures. Rats were euthanized by decapitation.

### Motor neuron isolation and purification

To generate primary motor neuron cultures, we adapted a published protocol intended for mice (*64*). Briefly, E14 embryos were removed from the womb and decapitated. The spinal cord of each embryo was removed from the body, cleaned of connective tissue, and cut into small pieces. Spinal cord pieces were incubated in 0.025% trypsin (15090046, Gibco) for 20 min. at 37°C. Dissociated cells were obtained by multiple trituration steps with 100 μL 4% bovine-serum albumin (A9418, Sigma-Aldrich) and 100 μL 1 mg/mL DNase (DN25, Sigma-Aldrich) diluted in 900 μL L-15 medium (11415049, Gibco), containing 3.5 mg/ml glucose (A2494001, Gibco), 10,000 U/mL Penicillin-Streptomycin (15140122, Gibco), 2% heat-inactivated horse serum (26050088, Gibco), 0.02 mM progesterone (P8783, Sigma-Aldrich), 0.01 mg/mL Insulin (I6634, Sigma-Aldrich), Putrescin (P5780, Sigma-Aldrich), Conalbumin (C7786, Sigma-Aldrich), and 0.001 mg/mL Sodium Selenite (S5261, Sigma-Aldrich). Motor neuron enrichment at this stage was 3-5%, as determined by immunofluorescence staining with Islet1+2 (39.4D5, DSHB). The dissociated cells underwent density gradient centrifugation (830 x g, 15 min., room temperature), using Optiprep (D1556; Sigma-Aldrich) at a final concentration of 9.5%, increasing the proportion of motor neurons to 30-40%.

### Co-culture experiments

Passage 0 MuSCs or FIBs were thawed, cultured in growth medium (C-23060; PromoCell) with 1% L-glutamine-penicillin-streptomycin solution (G6784; Sigma), 15% fetal bovine serum (FBS; ALB-S1810, Biowest) for 5 days, and then plated at 10,000 cells/cm^2^, in 24-well plates containing glass coverslips (day -1). The next day (day 0), motor neurons were added at 5,000 cells/cm^2^, and the medium was changed to a 1:1 mixture of differentiation medium (C-23260; PromoCell) with 1% L-glutamine-penicillin-streptomycin solution (G6784; Sigma) and Neurobasal medium (A3582901; Gibco) with 2% B27 supplement (A3582801; Gibco), 0.25% L-Glutamine (25030032; Thermo Scientific), 1 ng/ml Glial Derived Neurotrophic Factor (450-51; PeProTech), 1 ng/ml Brain Derived Neurotrophic Factor (450-02; PeProTech) and 10 ng/ml Ciliary Derived Neurotrophic Factor (450-50; PeProTech)). After 24 hours (day 1), experiments were stopped, and the cells were processed for immunofluorescence and RNA extraction.

### Conditioned medium experiments

MuSCs or FIBs were plated at 5,000 cells/cm^2^ in 12-well plates containing glass coverslips, and grown in growth medium (C-23060; PromoCell) with 1% L-glutamine-penicillin-streptomycin solution (G6784; Sigma) and 15% fetal bovine serum (FBS; ALB-S1810, Biowest) for 3 days. Cells were then washed, and the medium changed to differentiation medium (C-23260; PromoCell) with 1% L-glutamine-penicillin-streptomycin solution (G6784; Sigma), for 4 days, with a medium change after 2 days. Conditioned medium was collected at day 5 (2 days of differentiation) and 7 (4 days of differentiation), and frozen at -80°C. Morphometric and molecular characteristics of these cells are presented elsewhere (*37*). Motor neurons were plated at 5,000 cells/cm^2^ in 24-well plates on glass coverslips, coated with 10 μg/ml Poly-L-Ornithine (P8638; Sigma-Aldrich) and 3 μg/ml Laminin (L2020; Sigma-Aldrich). The conditioned medium from both day 5 and 7 were thawed, pooled, and then centrifuged at 10.000 x g for 5 min at 4°C, to remove cell debris. The neurons were plated in 60% Neurobasal medium (described under co-culture experiments) and 40% conditioned medium. After 24 (day 1) and 48 (day 2) hours, experiments were stopped and processed for immunofluorescence and RNA extraction.

### Immunofluorescence, microscopy, and image analyses

Embedded muscle samples were cryosectioned, stained with primary mouse anti-dystrophin (D8168; Sigma-Aldrich) and a secondary fluorescent (568 nm) goat anti-mouse antibody (A-21144; Invitrogen), and mounted with coverglasses using mounting medium (P36931; Thermo Fisher Scientific) containing 4’,6-diamidino-2-phenylindole (DAPI). Samples were imaged with a 10×/0.30 NA objective and a 0.5× camera (DP71, Olympus) mounted on a BX51 Olympus microscope, and analyzed using a semi-automated macro built in Fiji (*65*) (v. 1.51) (*66*), for muscle fiber cross-sectional area.

Cells were fixed using Histofix (Histolab), and then stained for ChAT (AB144P; Sigma-Aldrich) and Tau1 (GTX130462; GeneTex) with appropriate fluorescent secondary antibodies (A-11057, Invitrogen and 711-545-152, Jackson ImmunoResearch Laboratories). Coverslips were mounted on slides with mounting medium (P36931; Thermo Fisher Scientific) containing DAPI. For the co-cultures, four images were taken at fixed spots (north, south, east, west), with a 10×/0.30 NA objective and a 0.5× camera (DP71, Olympus) mounted on a BX51 Olympus microscope. The images were analyzed in Fiji, using the ObjectJ and SNT (*67*) plugins, for number of motor neurons and neurites per area, as well as neurite length per area. For the motor neuron cultures, entire coverslips were imaged with an AxioScan.Z1 slide scanner (Carl Zeiss) using a plan-apochromat 10×/0.45 NA objective and a MultiBand filter cube (DAPI/FITC/TexasRed) with excitation wavelengths of 353, 493 and 577 nm and both coarse and fine focusing steps. Channels were imaged sequentially with an AxioCam MR R3 and a 10% overlap between images. Merged images were stitched using ZEN blue software (Carl Zeiss) and analyzed using a semi-automated macro in Fiji. In brief, the DAPI channel was used for segmenting out nuclei using a manually selected threshold for each image. Manual corrections were then made to split clumps of nuclei or add missing nuclei. Subsequently, the intensity of the ChAT and Tau1 signal within each nuclei region was determined and the total area covered by Tau1 signal was measured, using a manual threshold. Intensity histograms were used to manually define nuclei positive for ChAT and Tau1 and their number per mm^2^ was calculated.

### Serum CAF ELISA

Serum CAF concentration was measured by ELISA following the manufactures instructions (ab216945, Abcam, Cambridge, United Kingdom). Optical density was recorded at 450 nm using a microplate ELISA reader (Multiscan FC, Thermo-Fisher Scientific). Biological replicates were analyzed in duplicate, and the mean of each duplicate was used in statistical analysis.

### RNA extraction

Total RNA was extracted from the cell cultures with TriReagent (TR118; Molecular Research) as described in Bechshøft et al. (*37*). For the conditioned medium experiment 400 ng total RNA purified using TriReagent from baker’s yeast (Frisk Bagegær, De Danske Gærfabrikker, Grenaa, Denmark) was added to the TriReagent to minimize loss of rat neuron RNA due to the very low amount. For the Co-culture experiment, the human RNA serves as protective carrier for the rat RNA (yield MuSC: 323±177, Fib: 752±346 ng total RNA).

### RNAseq

RNAseq was performed by a commercial company (Azenta, Liepzig, Germany). Briefly, RNA samples were quantified using Qubit 4.0 Fluorometer (Life Technologies, Carlsbad, CA, USA) and RNA integrity was checked with RNA Kit on Agilent 5300 Fragment Analyzer (Agilent Technologies, Palo Alto, CA, USA).

RNA sequencing libraries were prepared using the NEBNext Ultra II RNA Library Prep Kit for Illumina following manufacturer’s instructions (NEB, Ipswich, MA, USA). Briefly, mRNAs were first enriched with Oligo(dT) beads. Enriched mRNAs were fragmented according to manufacturer’s instruction. First strand and second strand cDNAs were subsequently synthesized. cDNA fragments were end repaired and adenylated at 3’ends, and universal adapters were ligated to cDNA fragments, followed by index addition and library enrichment by limited-cycle PCR. Sequencing libraries were validated using NGS Kit on the Agilent 5300 Fragment Analyzer (Agilent Technologies, Palo Alto, CA, USA), and quantified by using Qubit 4.0 Fluorometer (Invitrogen, Carlsbad, CA).

The sequencing libraries were multiplexed and loaded on the flow cell on the Illumina NovaSeq 6000 instrument according to manufacturer’s instructions. The samples were sequenced using a 2x150 Pair-End (PE) configuration v1.5. Image analysis and base calling were conducted by the NovaSeq Control Software v1.7 on the NovaSeq instrument. Raw sequence data (.bcl files) generated from Illumina NovaSeq was converted into fastq files and de-multiplexed using Illumina bcl2fastq program version 2.20. One mismatch was allowed for index sequence identification. 17-77 million paired reads were obtained per sample.

### Bioinformatics

The reads were split into Human, Rat and Yeast reads using bbsplit (http://sourceforge.net/projects/bbmap/) by comparison to the Rat mRatBN7 and Human GRCh38 or Yeast R64 genomes. Ambiguous reads were excluded. Human and Rat reads were aligned to the respective genomes and transcripts (exons) counted using SubRead v2.0.3 (*68*) resulting in 10−46, 0.6-6.0 or 0.1-0.8 million counts per sample, for human Co-culture, rat Co-culture and rat Conditioned, respectively.

Preliminary inspection of marker genes suggested that some cell purifications had failed. Therefore, CDSeqR v1.0.9 (*69*) was use for deconvolution of the human samples into cell types. Four of the fibroblast cultures were confirmed by the expression pattern of myoblast/fibroblast markers to consist mainly of myoblasts. Also, the expression pattern (Human and Rat) of one myoblast sample was strongly deviating from all the other samples, with very low proliferation (MKI67/Mki67) and very high metabolism (RPL23A/Rpl23a, GAPDH/Gapdh and MT-CO1/Mt-co1). These five samples were therefore excluded from the remaining analyses.

Count normalization and differential analyses were performed using DESeq2 v1.34.0 (*70*). For MuSC versus FIB, subjects were included as random factors and Age as fixed for the conditioned medium experiment. To account for log-fold inflation of the low counts, the lcfshrink method of DESeq2 was used. GO-term enrichment analysis was performed using topGO v2.52.0 (doi:10.18129/B9.bioc.topGO) with the elim algorithm and fisher test.

### Statistical Analysis

Data are shown in tables as mean ± SEM unless specified otherwise. RNA sequencing data are shown as DESeq2 normalized counts or log2 fold changes. Statistical analyses were conducted in GraphPad Prism (v.10, GraphPad Software) or DESeq2 (v1.34.0). The statistical test used for each dataset is provided in the accompanying figure legend. P values of <0.05 were considered significant.

## Funding

We gratefully acknowledge funding from:

The Lundbeck Foundation (R344-2020-254, R402-2022-1387) (C.S. and A.L.M).

Nordea Foundation (Centre for Healthy Aging) (M.K.).

Danielsen Fond (C.S).

Association pour le Développement de la Neurogénétique (A.J).

AFM Téléthon through the MyoNeurALP strategic grant (A.J., L.S. and B.C).

## Author contributions

CS, PS, AJ, LS, BC, and ALM conceived and designed research.

CS, PS, and RBS performed experiments.

CS, PS, and ALM analyzed data.

CS, PS, RBS, MK, AJ, LS, BC, and ALM interpreted results of experiments.

CS and PS prepared figures.

CS, PS, and ALM drafted manuscript.

All authors edited, revised, and approved the final version of the manuscript.

## Data and materials availability

All data are available upon request to the corresponding author.

## Supplemental material legends

**Fig. S1**: Differentially regulated human genes in Young vs Old after 1 day for FIB (blue) and MuSC (red). Data are means (horizontal line) with individual values. Human n: Old: 12 (MuSC) and 8 (FIB). Young: 8 (MuSC) and 6 (FIB). Rat n: 7. Data were analyzed with DESeq2, see bioinformatics section. Abbreviations: MuSC, muscle stem cell; FIB, muscle fibroblast.

**Table S1**: Participants characteristics. Age (years), height (cm), weight (kg) and BMI (kg/m^2^) for all participants. Data are means ± SD with ranges. BMI, body mass index.

## References

1. B. E. Tomlinson, D. Irving, The numbers of limb motor neurons in the human lumbosacral cord throughout life. J. Neurol. Sci. 34, 213–219 (1977).

2. C. J. McNeil, T. J. Doherty, D. W. Stashuk, C. L. Rice, Motor unit number estimates in the tibialis anterior muscle of young, old, and very old men. Muscle & Nerve 31, 461–467 (2005).

3. C. Soendenbroe, J. L. Andersen, A. L. Mackey, Muscle-nerve communication and the molecular assessment of human skeletal muscle denervation with aging. American Journal of Physiology-Cell Physiology 321, C317–C329 (2021).

4. J. S. McPhee, J. Cameron, T. Maden-Wilkinson, M. Piasecki, M. H. Yap, D. A. Jones, H. Degens, The Contributions of Fiber Atrophy, Fiber Loss, In Situ Specific Force, and Voluntary Activation to Weakness in Sarcopenia. J. Gerontol. A Biol. Sci. Med. Sci. 73, 1287–1294 (2018).

5. J. Lexell, C. C. Taylor, M. Sjöström, What is the cause of the ageing atrophy? Total number, size and proportion of different fiber types studied in whole vastus lateralis muscle from 15- to 83-year-old men. J. Neurol. Sci. 84, 275–294 (1988).

6. L. Madaro, M. Passafaro, D. Sala, U. Etxaniz, F. Lugarini, D. Proietti, M. V. Alfonsi, C. Nicoletti, S. Gatto, M. De Bardi, R. Rojas-García, L. Giordani, S. Marinelli, V. Pagliarini, C. Sette, A. Sacco, P. L. Puri, Denervation-activated STAT3-IL-6 signalling in fibro-adipogenic progenitors promotes myofibres atrophy and fibrosis. Nat Cell Biol 20, 917–927 (2018).

7. D. L. Rebolledo, D. González, J. Faundez-Contreras, O. Contreras, C. P. Vio, J. E. Murphy-Ullrich, K. E. Lipson, E. Brandan, Denervation-induced skeletal muscle fibrosis is mediated by CTGF/CCN2 independently of TGF-β. Matrix Biol 82, 20–37 (2019).

8. C. L. Gatchalian, M. Schachner, J. R. Sanes, Fibroblasts that proliferate near denervated synaptic sites in skeletal muscle synthesize the adhesive molecules tenascin(J1), N-CAM, fibronectin, and a heparan sulfate proteoglycan. J Cell Biol 108, 1873–1890 (1989).

9. M. J. Petrany, C. O. Swoboda, C. Sun, K. Chetal, X. Chen, M. T. Weirauch, N. Salomonis, D. P. Millay, Single-nucleus RNA-seq identifies transcriptional heterogeneity in multinucleated skeletal myofibers. Nat Commun 11, 6374 (2020).

10. M. N. Wosczyna, T. A. Rando, A Muscle Stem Cell Support Group: Coordinated Cellular Responses in Muscle Regeneration. Dev Cell 46, 135–143 (2018).

11. M. M. Murphy, J. A. Lawson, S. J. Mathew, D. A. Hutcheson, G. Kardon, Satellite cells, connective tissue fibroblasts and their interactions are crucial for muscle regeneration. Development 138, 3625–3637 (2011).

12. E. A. Connor, U. J. McMahan, Cell accumulation in the junctional region of denervated muscle. J Cell Biol 104, 109–120 (1987).

13. C. Nicoletti, X. Wei, U. Etxaniz, C. D’Ercole, L. Madaro, R. Perera, P. L. Puri, Muscle denervation promotes functional interactions between glial and mesenchymal cells through NGFR and NGF. iScience 26, 107114 (2023).

14. M. Theret, F. M. V. Rossi, O. Contreras, Evolving Roles of Muscle-Resident Fibro-Adipogenic Progenitors in Health, Regeneration, Neuromuscular Disorders, and Aging. Front Physiol 12, 673404 (2021).

15. A. Uezumi, M. Ikemoto-Uezumi, H. Zhou, T. Kurosawa, Y. Yoshimoto, M. Nakatani, K. Hitachi, H. Yamaguchi, S. Wakatsuki, T. Araki, M. Morita, H. Yamada, M. Toyoda, N. Kanazawa, T. Nakazawa, J. Hino, S.-I. Fukada, K. Tsuchida, Mesenchymal Bmp3b expression maintains skeletal muscle integrity and decreases in age-related sarcopenia. J Clin Invest 131, e139617, 139617 (2021).

16. C. Lepper, T. A. Partridge, C.-M. Fan, An absolute requirement for Pax7-positive satellite cells in acute injury-induced skeletal muscle regeneration. Development 138, 3639–3646 (2011).

17. E. Barruet, S. M. Garcia, K. Striedinger, J. Wu, S. Lee, L. Byrnes, A. Wong, S. Xuefeng, S. Tamaki, A. S. Brack, J. H. Pomerantz, Functionally heterogeneous human satellite cells identified by single cell RNA sequencing. Elife 9, e51576 (2020).

18. K. A. Murach, C. S. Fry, E. E. Dupont-Versteegden, J. J. McCarthy, C. A. Peterson, Fusion and beyond: Satellite cell contributions to loading-induced skeletal muscle adaptation. FASEB J 35, e21893 (2021).

19. W. Liu, A. Klose, S. Forman, N. D. Paris, L. Wei-LaPierre, M. Cortés-Lopéz, A. Tan, M. Flaherty, P. Miura, R. T. Dirksen, J. V. Chakkalakal, Loss of adult skeletal muscle stem cells drives age-related neuromuscular junction degeneration. eLife 6, e26464 (2017).

20. R. Mills, H. Taylor-Weiner, J. C. Correia, L. Z. Agudelo, I. Allodi, C. Kolonelou, V. Martinez-Redondo, D. M. S. Ferreira, S. Nichterwitz, L. H. Comley, V. Lundin, E. Hedlund, J. L. Ruas, A. I. Teixeira, Neurturin is a PGC-1α1-controlled myokine that promotes motor neuron recruitment and neuromuscular junction formation. Mol Metab 7, 12–22 (2018).

21. J. V. Montoya G, J. J. Sutachan, W. S. Chan, A. Sideris, T. J. J. Blanck, E. Recio-Pinto, Muscle-conditioned media and cAMP promote survival and neurite outgrowth of adult spinal cord motor neurons. Exp Neurol 220, 303–315 (2009).

22. R. J. Chai, J. Vukovic, S. Dunlop, M. D. Grounds, T. Shavlakadze, Striking denervation of neuromuscular junctions without lumbar motoneuron loss in geriatric mouse muscle. PLoS ONE 6, e28090 (2011).

23. R. W. Castro, M. C. Lopes, R. E. Settlage, G. Valdez, Aging alters mechanisms underlying voluntary movements in spinal motor neurons of mice, primates, and humans. JCI Insight 8, e168448 (2023).

24. M. R. Deschenes, Motor unit and neuromuscular junction remodeling with aging. Curr Aging Sci 4, 209–220 (2011).

25. E. J. Jones, J. Piasecki, A. Ireland, D. W. Stashuk, P. J. Atherton, B. E. Phillips, J. S. McPhee, M. Piasecki, Lifelong exercise is associated with more homogeneous motor unit potential features across deep and superficial areas of vastus lateralis. Geroscience 43, 1555–1565 (2021).

26. L. M. Snow, L. K. Mcloon, L. V. Thompson, Adult and developmental myosin heavy chain isoforms in soleus muscle of aging Fischer Brown Norway rat. *The Anatomical Record Part A: Discoveries in Molecular*, Cellular, and Evolutionary Biology 286A, 866–873 (2005).

27. R. T. Hepple, C. L. Rice, Innervation and neuromuscular control in ageing skeletal muscle. J. Physiol. (Lond.) 594, 1965–1978 (2016).

28. G. A. Power, B. H. Dalton, D. G. Behm, A. A. Vandervoort, T. J. Doherty, C. L. Rice, Motor Unit Number Estimates in Masters Runners: Use It or Lose It? Medicine & Science in Sports & Exercise 42, 1644–1650 (2010).

29. V. Sonjak, K. Jacob, J. A. Morais, M. Rivera-Zengotita, S. Spendiff, C. Spake, T. Taivassalo, S. Chevalier, R. T. Hepple, Fidelity of muscle fibre reinnervation modulates ageing muscle impact in elderly women. The Journal of Physiology 597, 5009–5023 (2019).

30. C. Soendenbroe, M. F. Flindt Heisterberg, P. Schjerling, M. Kjaer, J. L. Andersen, A. L. Mackey, Human skeletal muscle acetylcholine receptor gene expression in elderly males performing heavy resistance exercise. Am J Physiol Cell Physiol, doi: 10.1152/ajpcell.00365.2021 (2022).

31. T. Mars, K. J. Yu, X. M. Tang, A. F. Miranda, Z. Grubic, F. Cambi, M. P. King, Differentiation of glial cells and motor neurons during the formation of neuromuscular junctions in cocultures of rat spinal cord explant and human muscle. J Comp Neurol 438, 239–251 (2001).

32. C. C. Agley, A. M. Rowlerson, C. P. Velloso, N. R. Lazarus, S. D. R. Harridge, Human skeletal muscle fibroblasts, but not myogenic cells, readily undergo adipogenic differentiation. J. Cell. Sci. 126, 5610–5625 (2013).

33. E. Negroni, M. Kondili, L. Muraine, M. Bensalah, G. S. Butler-Browne, V. Mouly, A. Bigot, C. Trollet, Muscle fibro-adipogenic progenitors from a single-cell perspective: Focus on their “virtual” secretome. Front Cell Dev Biol 10, 952041 (2022).

34. A. Karlsen, C.-Y. C. Yeung, P. Schjerling, L. Denz, C. Hoegsbjerg, J. R. Jakobsen, M. R. Krogsgaard, M. Koch, S. Schiaffino, M. Kjaer, A. L. Mackey, Distinct myofibre domains of the human myotendinous junction revealed by single nucleus RNA-seq. J Cell Sci, jcs.260913 (2023).

35. J. Delile, T. Rayon, M. Melchionda, A. Edwards, J. Briscoe, A. Sagner, Single cell transcriptomics reveals spatial and temporal dynamics of gene expression in the developing mouse spinal cord. Development 146, dev173807 (2019).

36. J. A. Blum, S. Klemm, J. L. Shadrach, K. A. Guttenplan, L. Nakayama, A. Kathiria, P. T. Hoang, O. Gautier, J. A. Kaltschmidt, W. J. Greenleaf, A. D. Gitler, Single-cell transcriptomic analysis of the adult mouse spinal cord reveals molecular diversity of autonomic and skeletal motor neurons. Nat Neurosci 24, 572–583 (2021).

37. C. J. L. Bechshøft, S. M. Jensen, P. Schjerling, J. L. Andersen, R. B. Svensson, C. S. Eriksen, N. S. Mkumbuzi, M. Kjaer, A. L. Mackey, Age and prior exercise in vivo determine the subsequent in vitro molecular profile of myoblasts and nonmyogenic cells derived from human skeletal muscle. Am. J. Physiol., Cell Physiol. 316, C898–C912 (2019).

38. T. Wang, J. Chen, C.-X. Tang, X.-Y. Zhou, D.-S. Gao, Inverse Expression Levels of EphrinA3 and EphrinA5 Contribute to Dopaminergic Differentiation of Human SH-SY5Y Cells. J Mol Neurosci 59, 483–492 (2016).

39. S. Negro, F. Lessi, E. Duregotti, P. Aretini, M. La Ferla, S. Franceschi, M. Menicagli, E. Bergamin, E. Radice, M. Thelen, A. Megighian, M. Pirazzini, C. M. Mazzanti, M. Rigoni, C. Montecucco, CXCL12α/SDF-1 from perisynaptic Schwann cells promotes regeneration of injured motor axon terminals. EMBO Mol Med 9, 1000–1010 (2017).

40. X. Caubit, M. Thoby-Brisson, N. Voituron, P. Filippi, M. Bévengut, H. Faralli, S. Zanella, G. Fortin, G. Hilaire, L. Fasano, Teashirt 3 regulates development of neurons involved in both respiratory rhythm and airflow control. J Neurosci 30, 9465–9476 (2010).

41. M. Gao, Q. Dong, Z. Yang, D. Zou, Y. Han, Z. Chen, R. Xu, Long non-coding RNA H19 regulates neurogenesis of induced neural stem cells in a mouse model of closed head injury. Neural Regen Res 19, 872–880 (2024).

42. Y. Sun, Y. Jin, An intraflagellar transport dependent negative feedback regulates the MAPKKK DLK-1 to protect cilia from degeneration. Proc Natl Acad Sci U S A 120, e2302801120 (2023).

43. C. Soendenbroe, C. L. Dahl, C. Meulengracht, M. Tamáš, R. B. Svensson, P. Schjerling, M. Kjaer, J. L. Andersen, A. L. Mackey, Preserved stem cell content and innervation profile of elderly human skeletal muscle with lifelong recreational exercise. The Journal of Physiology 600, 1969–1989 (2022).

44. M. N. Wosczyna, C. T. Konishi, E. E. Perez Carbajal, T. T. Wang, R. A. Walsh, Q. Gan, M. W. Wagner, T. A. Rando, Mesenchymal Stromal Cells Are Required for Regeneration and Homeostatic Maintenance of Skeletal Muscle. Cell Rep 27, 2029–2035.e5 (2019).

45. K. Wang, O. K. Yaghi, R. G. Spallanzani, X. Chen, D. Zemmour, N. Lai, I. M. Chiu, C. Benoist, D. Mathis, Neuronal, stromal, and T-regulatory cell crosstalk in murine skeletal muscle. Proc Natl Acad Sci U S A 117, 5402–5408 (2020).

46. K. Mis, Z. Grubic, P. Lorenzon, M. Sciancalepore, T. Mars, S. Pirkmajer, In Vitro Innervation as an Experimental Model to Study the Expression and Functions of Acetylcholinesterase and Agrin in Human Skeletal Muscle. Molecules 22, E1418 (2017).

47. A. Barik, L. Li, A. Sathyamurthy, W.-C. Xiong, L. Mei, Schwann Cells in Neuromuscular Junction Formation and Maintenance. J Neurosci 36, 9770–9781 (2016).

48. J. V. Chakkalakal, H. Nishimune, J. L. Ruas, B. M. Spiegelman, J. R. Sanes, Retrograde influence of muscle fibers on their innervation revealed by a novel marker for slow motoneurons. Development 137, 3489–3499 (2010).

49. B. K. Pedersen, Physical activity and muscle–brain crosstalk. Nat Rev Endocrinol 15, 383–392 (2019).

50. A. Florin, C. Lambert, C. Sanchez, J. Zappia, N. Durieux, A. M. Tieppo, A. Mobasheri, Y. Henrotin, The secretome of skeletal muscle cells: A systematic review. Osteoarthritis and Cartilage Open 2, 100019 (2020).

51. E. E. Zahavi, A. Ionescu, S. Gluska, T. Gradus, K. Ben-Yaakov, E. Perlson, A compartmentalized microfluidic neuromuscular co-culture system reveals spatial aspects of GDNF functions. J. Cell. Sci. 128, 1241–1252 (2015).

52. L. Le Gall, W. J. Duddy, C. Martinat, V. Mariot, O. Connolly, V. Milla, E. Anakor, Z. G. Ouandaogo, S. Millecamps, J. Lainé, U. G. Vijayakumar, S. Knoblach, C. Raoul, O. Lucas, J. P. Loeffler, P. Bede, A. Behin, H. Blasco, G. Bruneteau, M. Del Mar Amador, D. Devos, A. Henriques, A. Hesters, L. Lacomblez, P. Laforet, T. Langlet, P. Leblanc, N. Le Forestier, T. Maisonobe, V. Meininger, L. Robelin, F. Salachas, T. Stojkovic, G. Querin, J. Dumonceaux, G. Butler Browne, J.-L. González De Aguilar, S. Duguez, P. F. Pradat, Muscle cells of sporadic amyotrophic lateral sclerosis patients secrete neurotoxic vesicles. J Cachexia Sarcopenia Muscle, doi: 10.1002/jcsm.12945 (2022).

53. C. E. Stewart, A. P. Sharples, Aging, Skeletal Muscle, and Epigenetics. Plast Reconstr Surg 150, 27S–33S (2022).

54. J. Lund, S. A. Helle, Y. Li, N. G. Løvsletten, H. K. Stadheim, J. Jensen, E. T. Kase, G. H. Thoresen, A. C. Rustan, Higher lipid turnover and oxidation in cultured human myotubes from athletic versus sedentary young male subjects. Sci Rep 8, 17549 (2018).

55. A. Blasco, S. Gras, G. Mòdol-Caballero, O. Tarabal, A. Casanovas, L. Piedrafita, A. Barranco, T. Das, S. L. Pereira, X. Navarro, R. Rueda, J. E. Esquerda, J. Calderó, Motoneuron deafferentation and gliosis occur in association with neuromuscular regressive changes during ageing in mice. J Cachexia Sarcopenia Muscle 11, 1628–1660 (2020).

56. Y. Kawamura, H. Okazaki, P. C. O’Brien, P. J. Dych, Lumbar motoneurons of man: I) number and diameter histogram of alpha and gamma axons of ventral root. J Neuropathol Exp Neurol 36, 853–860 (1977).

57. K. R. Mittal, F. H. Logmani, Age-related reduction in 8th cervical ventral nerve root myelinated fiber diameters and numbers in man. J Gerontol 42, 8–10 (1987).

58. C. J. Lukasiewicz, G. J. Tranah, D. S. Evans, P. M. Coen, H. N. Barnes, Z. Huo, K. A. Esser, X. Zhang, C. Wolff, K. Wu, N. E. Lane, S. B. Kritchevsky, A. B. Newman, S. R. Cummings, P. M. Cawthon, R. T. Hepple, Higher expression of denervation-responsive genes is negatively associated with muscle volume and performance traits in the study of muscle, mobility, and aging (SOMMA). Aging Cell, e14115 (2024).

59. U. R. Mikkelsen, C. Couppé, A. Karlsen, J. F. Grosset, P. Schjerling, A. L. Mackey, H. H. Klausen, S. P. Magnusson, M. Kjær, Life-long endurance exercise in humans: circulating levels of inflammatory markers and leg muscle size. Mech. Ageing Dev. 134, 531–540 (2013).

60. M. Piasecki, A. Ireland, J. Piasecki, D. W. Stashuk, A. Swiecicka, M. K. Rutter, D. A. Jones, J. S. McPhee, Failure to expand the motor unit size to compensate for declining motor unit numbers distinguishes sarcopenic from non-sarcopenic older men. J. Physiol. (Lond.) 596, 1627–1637 (2018).

61. N. J. Pillon, B. M. Gabriel, L. Dollet, J. A. B. Smith, L. Sardón Puig, J. Botella, D. J. Bishop, A. Krook, J. R. Zierath, Transcriptomic profiling of skeletal muscle adaptations to exercise and inactivity. Nat Commun 11, 470 (2020).

62. A. B. Moy, A. Kamath, S. Ternes, J. Kamath, The Challenges to Advancing Induced Pluripotent Stem Cell-Dependent Cell Replacement Therapy. Med Res Arch 11, 4784 (2023).

63. J. Bergstrom, Percutaneous needle biopsy of skeletal muscle in physiological and clinical research. Scand. J. Clin. Lab. Invest. 35, 609–616 (1975).

64. A. Jacquier, V. Risson, L. Schaeffer, Modeling Charcot-Marie-Tooth Disease In Vitro by Transfecting Mouse Primary Motoneurons. JoVE (Journal of Visualized Experiments), e57988 (2019).

65. J. Schindelin, I. Arganda-Carreras, E. Frise, V. Kaynig, M. Longair, T. Pietzsch, S. Preibisch, C. Rueden, S. Saalfeld, B. Schmid, J.-Y. Tinevez, D. J. White, V. Hartenstein, K. Eliceiri, P. Tomancak, A. Cardona, Fiji: an open-source platform for biological-image analysis. Nature Methods 9, 676–682 (2012).

66. A. Karlsen, R. L. Bechshøft, N. M. Malmgaard-Clausen, J. L. Andersen, P. Schjerling, M. Kjaer, A. L. Mackey, Lack of muscle fibre hypertrophy, myonuclear addition, and satellite cell pool expansion with resistance training in 83-94-year-old men and women. Acta Physiol (Oxf) 227, e13271 (2019).

67. C. Arshadi, U. Günther, M. Eddison, K. I. S. Harrington, T. A. Ferreira, SNT: a unifying toolbox for quantification of neuronal anatomy. Nat Methods 18, 374–377 (2021).

68. Y. Liao, G. K. Smyth, W. Shi, The Subread aligner: fast, accurate and scalable read mapping by seed-and-vote. Nucleic Acids Res 41, e108 (2013).

69. K. Kang, C. Huang, Y. Li, D. M. Umbach, L. Li, CDSeqR: fast complete deconvolution for gene expression data from bulk tissues. BMC Bioinformatics 22, 262 (2021).

70. M. I. Love, W. Huber, S. Anders, Moderated estimation of fold change and dispersion for RNA-seq data with DESeq2. Genome Biol 15, 550 (2014).

