## Supplemental material for "Muscle Fibroblasts and Stem Cells Stimulate Motor Neurons in An Age and Exercise-Dependent Manner"

### Supplemental figure S1

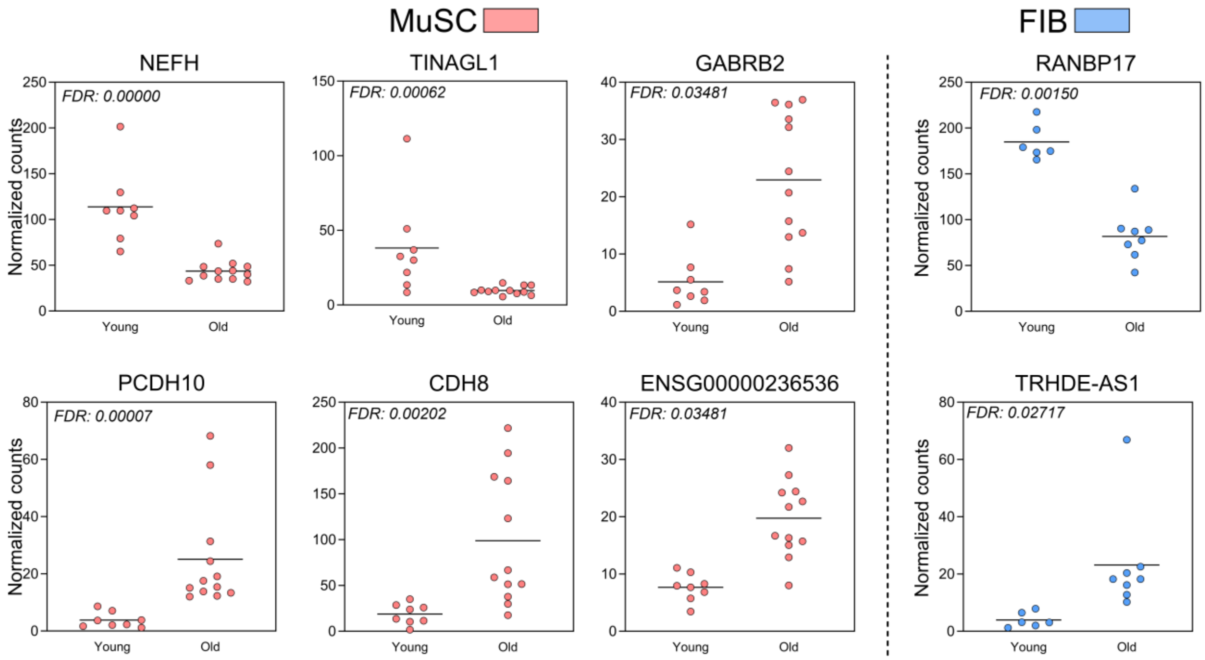

### Supplemental table 1

| | Young (n=21) | | | | Old LLEX (n=7) | | Old SED (n=17) | | | |
|---|---|---|---|---|---|---|---|---|---|---|
| | Female (n=12) | | Male (n=9) | | Male (n=7) | | Female (n=11) | | Male (n=6) | |
| | *Mean SD* | *min maks* | *Mean SD* | *min maks* | *Mean SD* | *min maks* | *Mean SD* | *min maks* | *Mean SD* | *min maks* |
| Age | 23 ± 3 | 20 - 28 | 26 ± 5 | 20 - 34 | 72 ± 2 | 69 - 76 | 74 ± 3 | 71 - 78 | 73 ± 4 | 68 - 78 |
| Height (cm) | 168 ± 7 | 157 - 177 | 181 ± 8 | 169 - 191 | 176 ± 6 | 168 - 182 | 166 ± 3 | 162 - 169 | 176 ± 9 | 161 - 188 |
| Weight (kg) | 64 ± 8 | 53 - 75 | 81 ± 15 | 62 - 105 | 76 ± 9 | 70 - 94 | 69 ± 10 | 57 - 84 | 82 ± 6 | 71 - 88 |
| BMI (kg/m$^2$) | 23 ± 2 | 19 - 26 | 25 ± 3 | 20 - 30 | 25 ± 3 | 22 - 30 | 25 ± 4 | 20 - 30 | 27 ± 3 | 23 - 32 |
